# Single-cell secretion analysis reveals a dual role for IL-10 in restraining and resolving the TLR4-induced inflammatory response

**DOI:** 10.1101/2020.10.22.351254

**Authors:** Amanda F. Alexander, Hannah Forbes, Kathryn Miller-Jensen

## Abstract

Following TLR4 stimulation of macrophages, negative feedback mediated by the anti-inflammatory cytokine IL-10 limits the inflammatory response. However, extensive cell-to-cell variability in TLR4-stimulated cytokine secretion raises questions about how negative feedback is robustly implemented. To explore this, we characterized the TLR4-stimulated secretion program in primary murine macrophages using a single-cell microwell assay that enabled evaluation of functional autocrine IL-10 signaling. High-dimensional analysis of single-cell data revealed three distinct tiers of TLR4-induced proinflammatory activation based on levels of cytokine secretion. Surprisingly, while IL-10 inhibits TLR4-induced activation in the highest tier, it also contributes to the TLR4-induced activation threshold by regulating which cells transition from non-secreting to secreting states. This role for IL-10 in restraining TLR4 inflammatory activation is largely mediated by intermediate IFN-β signaling, while TNF-a likely mediates response resolution by IL-10. Thus, cell-to-cell variability in cytokine regulatory motifs provides a means to tailor the TLR4-induced inflammatory response.

## Introduction

In the innate immune system, over-response, hyperinflammation, and dysregulation can lead to tissue damage (e.g., sepsis) and cause various autoimmune and metabolic diseases (Medzhitov, 2008). Therefore, tight regulation of the inflammatory response is necessary to maintain immune health (Peter J. Murray & Smale, 2012). A critical regulatory component shaping the innate immune response is cell-to-cell communication. Macrophages and dendritic cells both secrete cytokines and chemokines during inflammation, and also respond to those extracellular signals (Lee et al., 2009; Xue et al., 2015). TLR4 stimulation results in the production of proinflammatory cytokines like tumor necrosis factor-a (TNF), immuno-regulatory cytokines like interferon-β (IFN-β), and anti-inflammatory cytokines like IL-10. These proteins and other secreted factors are used by innate immune cells in both positive and negative feedback motifs mediated through cell-to-cell communication (Gottschalk et al., 2019; Hu et al., 2008; Lee et al., 2009).

Specifically, IL-10 is a known, potent, negative regulator necessary for inflammatory control and resolution (Peter J. Murray & Smale, 2012; Spits & de Waal Malefyt, 1992). Dysregulated or deficient IL-10 production is associated with inflammatory and autoimmune pathologies like septic shock, colitis, and rheumatoid arthritis (Shankar Subramanian Iyer & Cheng, 2012; Katsikis et al., 1994; Kühn et al., 1993). IL-10 negative feedback restricts activation of antigen presenting cells and the adaptive immune response, and specifically in TLR signaling, suppresses transcription of proinflammatory cytokines and chemokines (J. Chang et al., 2009; Conaway et al., 2017; Dagvadorj et al., 2008; Mittal & Roche, 2015; P. J. Murray, 2005). While there is debate about the exact mechanisms behind IL-10-mediated inhibition, several studies cite STAT3-mediated suppression of NF-κB as a key contributor (DRIESSLER et al., 2004; Hovsepian et al., 2013; Hutchins et al., 2013). Various efforts have attempted to harness the anti-inflammatory capabilities of IL-10 to treat immune diseases characterized by hyperinflammation (Cannella et al., 1996; O’Garra et al., 2008). Unfortunately, these efforts have been largely unsuccessful (Saxena et al., 2015), illustrating that more work is needed to fully elucidate IL-10 activity and regulation within the immune response.

One reason IL-10 negative feedback may be difficult to use therapeutically is because of its complex interactions with other regulatory motifs. Specifically, TNF and IFN-β are both regulators of IL-10 production, as well as key paracrine signals shaping the overall macrophage inflammatory response. IFN-β is an autocrine and paracrine mediator necessary for robust IL-10 secretion, resulting in a time lag in TLR4-mediated IL-10 production (E. Y. Chang et al., 2007; Howes et al., 2016; S. S. Iyer et al., 2010). Additionally, paracrine feedback mediated by “precocious” IFN-β-producing cells has been implicated as a driver of cell-to-cell variation at the transcriptional level in dendritic cells (Shalek et al., 2014). TNF also promotes the secretion of IL-10, as well as other proinflammatory cytokines, following TLR4 stimulation (Caldwell et al., 2014; Muldoon et al., 2020). Analogous to IFN-β, TNF is secreted heterogeneously by macrophages, and TNF “super-secretors” appear to play a predominant role in amplifying IL-6 and IL-10 secretion in surrounding cells (Xue et al., 2015). The biological significance of having IL-10 regulated by two intermediate cytokines exhibiting high cell-to-cell variability is largely unknown, and it raises questions regarding the role of these paracrine signaling motifs in shaping the overall response.

To explore these questions, we investigated the role of IL-10 negative feedback in shaping the TLR4-induced macrophage secretion response. To account for the extensive cell-to-cell heterogeneity in extracellular signaling, we used a microwell assay that allows autocrine signaling without interference from signals from neighboring cells to measure multiplexed cytokine secretion in individual cells. We characterized the TLR4-stimulated response in primary murine bone marrow-derived macrophages (BMDMs) with and without IL-10 negative feedback. We found that activated macrophages lie along a trimodal, heterogeneous, pro-inflammatory axis, with IL-10 negative feedback modulating the distribution of cells within each activation tier. While IL-10 restrained activation of highly activated macrophages, its primary effect, mediated by relatively low levels of IL-10 secretion, was to raise the threshold of activation for low and non-responder cells. IL-10’s effect on the TLR4 activation threshold was largely mediated by IFN-β, and together these cytokines regulated the fraction of cells that mount a robust inflammatory response in a population. In contrast, positive feedback by TNF increased IL-10 levels in highly activated macrophages, which in turn resolved the response in those cells. Thus, our study disentangles IL-10 signaling to reveal two distinct functional roles in negatively regulating the TLR4 inflammatory response.

## Results

### IL-10 suppresses TLR4-induced pro-inflammatory activation in a dose-dependent manner

Most secreted proteins induced by TLR4 stimulation in macrophages, including TNF, IL-6, and IL-12, carry out proinflammatory functions, but secretion of the anti-inflammatory cytokine IL-10 also plays an important role in the microbial response (Peter J. Murray & Smale, 2012). TLR4-induced IL-10 negatively regulates the inflammatory response, but IL-10 negative feedback is complicated by the fact that IL-10 is induced by the same signals it regulates, specifically TNF and IFN-β (Fig. 1A). To characterize the role of anti-inflammatory IL-10 within the TLR4 response, we stimulated BMDMs in population with TLR4 agonist lipopolysaccharide (LPS) at 100 ng/mL. We collected cell supernatant at 0, 4, 8, 10, 12, and 24 hours post-stimulation, and quantified the concentration of TNF and IL-10 by ELISA. We found concurrent increases in both TNF and IL-10 secretion through 8 hours, when TNF concentration peaked in the population. After 8 hours, TNF concentration slowly decreased, while IL-10 continued to rise, reaching its maximum concentration at approximately 12 hours and remaining at that level through 24 hours (Fig. 1B). These results are consistent with the previously reported time lag in TLR4-induced IL-10, relative to proinflammatory cytokines (E. Y. Chang et al., 2007; Ernst et al., 2019; S. S. Iyer et al., 2010).

**Fig. 1.**
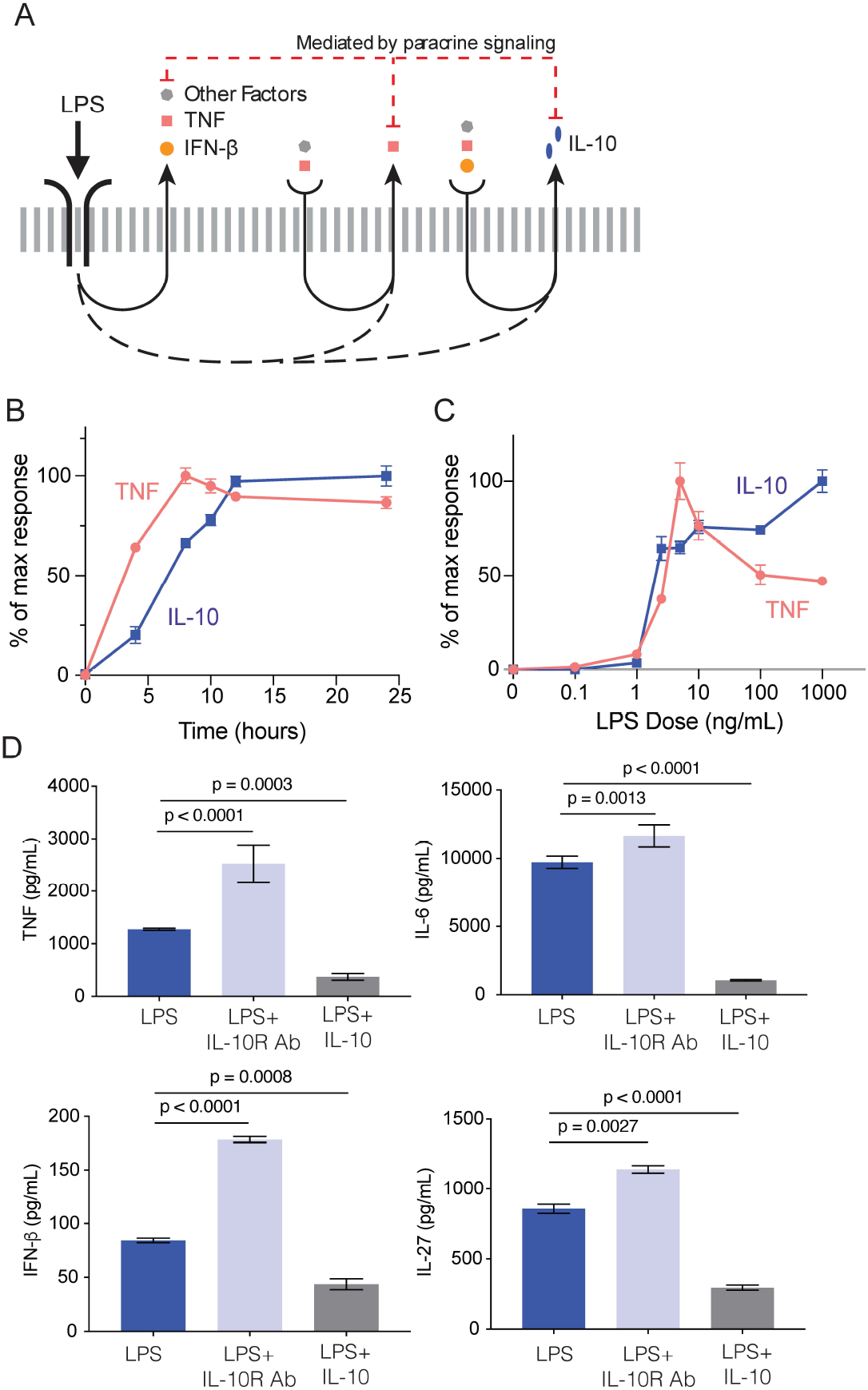
IL-10 suppresses TLR4-induced proinflammatory activation in a dose-dependent manner. (A) Schematic model illustrating how IL-10 negative feedback and paracrine signaling drive LPS-stimulated cytokine secretion. (B) BMDMs, derived from C57BL/6 mice, were stimulated with 100 ng/mL LPS for the indicated time points, after which cell supernatants were collected and TNF and IL-10 concentration measured by ELISA. (C) BMDMs were stimulated with the indicated doses of LPS for 8 hours, after which cell supernatants were collected and TNF and IL-10 concentration measured by ELISA. (D) BMDMs were stimulated with 100 ng/mL LPS alone, or co-stimulated with IL-10R Ab (30 μg/mL) to inhibit negative feedback, or co-stimulated with recombinant IL-10 (10 ng/mL) to enhance negative feedback. After 24 hours of stimulation cell supernatants were collected and cytokine production measured by ELISA. p-values determined by ordinary 2-way ANOVA. Data shown are mean ± SEM of 3 biological replicates.

To better understand the effect of stimulation dose on IL-10 secretion, we stimulated BMDMs in population with increasing doses of LPS spanning 5 orders of magnitude (0.1 – 1000 ng/mL). We collected cell supernatants after 8 and 24 hours, and measured TNF and IL-10 concentration by ELISA. The 8 and 24-hour time points were chosen because they marked peak and reduced TNF concentrations (Fig. 1B), and we hypothesized the reduction was caused at least in part by paracrine-mediated negative feedback from IL-10. The 8 and 24-hour dose responses both showed that above 1 ng/mL LPS there is steep induction of TNF and IL-10 secretion, with TNF reaching its peak between 5 and 10 ng/mL LPS (Fig. 1C and Fig. S1A).

This switch-like induction of TNF is consistent with a known threshold of TLR4-induced inflammatory activation mediated by MAPK intracellular signaling (Fig. S1B), which has been previously described (Gottschalk et al., 2016). We also found TNF secretion was significantly reduced following stimulation with high doses of LPS (100 and 1000 ng/mL). In contrast, IL-10 secretion continued to rise as LPS dose increased up to 1000 ng/mL at both 8 and 24 hours (Fig. 1C and Fig. S1A). These results reveal strong dose-dependence for IL-10 secretion and an inverse dose response for TNF and IL-10 at high LPS doses.

To directly measure how IL-10 negative feedback affects TLR4-induced proinflammatory secretion at high doses, we stimulated BMDMs with 100 ng/mL LPS alone, or in combination with an IL-10 receptor blocking antibody (IL-10R Ab) to remove negative feedback, or in combination with recombinant IL-10 to impose IL-10 negative feedback on the entire population. After 24 hours, cell supernatant was collected and several cytokines were measured by ELISA (Fig. 1D). We found that blocking IL-10 negative feedback via IL-10R Ab significantly increased the production of pro-inflammatory cytokines TNF, IL-6, IL-27, and IFN-β, consistent with previous reports (P. J. Murray, 2005; Xue et al., 2015). Consistently, co-stimulating BMDMs with LPS and recombinant IL-10 significantly decreased the secretion of TNF, IL-6, IL-27, and IFN-β. These data confirm that the anti-inflammatory effect of secreted IL-10 is both time and dose-sensitive.

### High-dimensional analysis reveals TLR4-activated macrophages exhibit dose-dependent digital and analog activation of the proinflammatory secretion program

Substantial cell-to-cell variability is a defining attribute of TLR-stimulated macrophage secretion (Lu et al., 2015; Ramji et al., 2019), and this variability plays a key role in permitting rapid and reproducible innate immune responses (Eberwine & Kim, 2015). While our secretion results from BMDMs in a population clearly show the potent anti-inflammatory effects of IL-10 signaling within the TLR4-induced response, they are unable to capture the cellular heterogeneity that exists. Therefore, to build upon our results at the population level and explore how IL-10 negative feedback varies across individual macrophages, we conducted multiplexed, single-cell, secretion profiling (Lu et al., 2013). We measured 12 cytokines, chemokines, and other canonical TLR4-induced secreted factors in order to sample the full range of macrophage functional responses (Table 1 and Materials and Methods). In this microwell format, cells are isolated such that functional autocrine signaling proceeds without interference from paracrine signals produced by neighboring cells.

**Table 1.** List of capture and detection antibody pairs used for single-cell secretion profiling.

| Antibody Pair | Vendor | Catalog No. |
| --- | --- | --- |
| TNF-α | Invitrogen | 88-7324-88 |
| IL-10 | R&D | DY417 |
| IL-6 | R&D | DY406 |
| IL-27 | R&D | DY1834 |
| CXCL1 | R&D | DY453 |
| CCL2 | R&D | DY479 |
| CCL3 | R&D | DY450 |
| CCL5 | R&D | DY478 |
| GM-CSF | R&D | DY415 |
| Chi3l3 | R&D | DY2446 |
| IFN-β | R&D | DY8234 |
| IL-12p40 | BD Biosciences | 555165 |

We first investigated the dose-dependence of IL-10 secretion in the TLR4 response in isolated cells. We stimulated BMDMs with a range of LPS doses (0, 10, 100, 1000 ng/mL) for 8 hours in the microwell device (Fig. 2A). As in the cell population, we observed dose-dependent increases in TLR4-mediated production of TNF and IL-10. Single-cell resolution demonstrated that this dose dependence was due to both the percentage of cells secreting each cytokine above background and the intensity of that secretion (Fig. 2B and 2C). Interestingly, the average observed fraction of cells secreting TNF was approximately 40%, while fewer than 20% of BMDMs secreted IL-10, even at the highest doses (Fig. 2C). We hypothesize that the fraction of cells responding in the device may be lower than in a population due to the loss of paracrine signaling in the microwell device, which has been shown to amplify cellular responses (Lee et al., 2009; Shalek et al., 2014; Xue et al., 2015; Youk & Lim, 2014).

**Fig. 2.**
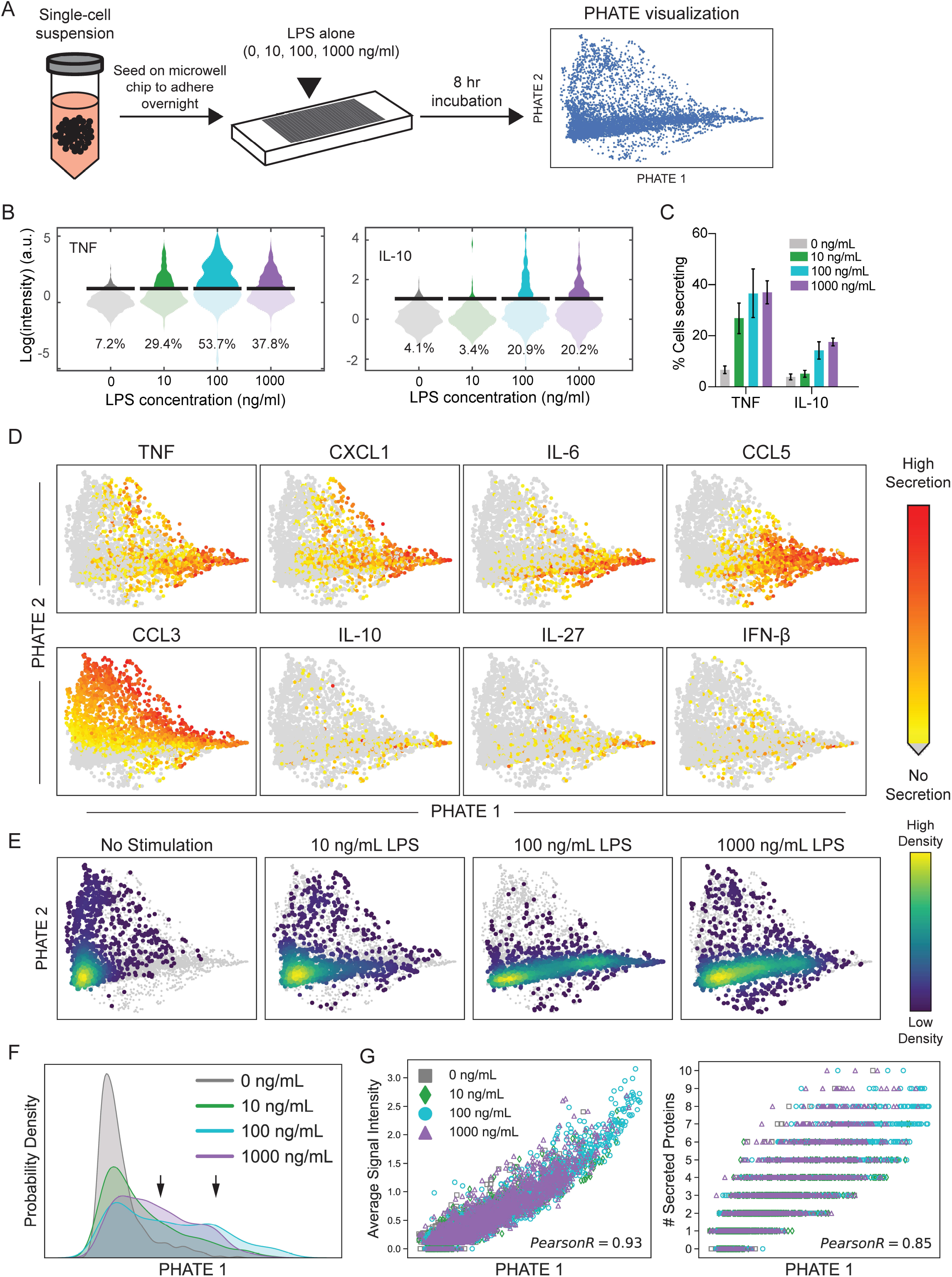
High-dimensional analysis reveals TLR4-activated macrophages exhibit dose-dependent digital and analog activation of the proinflammatory secretion program. (A) Experimental set-up for single-cell secretion profiling in the microwell device. BMDMs were seeded onto the PDMS microwell chip and allowed to adhere overnight. Individual cells were stimulated with increasing doses (0, 10, 100, 1000 ng/mL) of LPS for 8 hours. High-dimensional secretion data was then visualized by PHATE analysis. (B) Violin plots of single-cell secretion from individual BMDMs stimulated with increasing doses of LPS in the microwell assay. Black bar indicates fluorescent threshold of detection, arbitrary units (a.u.). Data are one representative experiment. (C) Bar graph depicts average percentage of cells secreting TNF and IL-10 above threshold in response to the indicated LPS stimulation dose after 8 hours in the microwell assay. Data shown are mean with 95% confidence intervals calculated by bootstrapping. (D and E) 2D PHATE embeddings of 10-dimensional single-cell secretion from isolated BMDMs stimulated with LPS as indicated in A. Data shown include activated macrophages (i.e. secreting at least 1 cytokine from Table 1) colored by (D) relative intensity of the indicated cytokine or (E) the kernel density estimate for cells at the indicated LPS stimulation dose (other doses shown in grey for visualization). Data pooled from 3 independent experiments. (F) Kernel density estimation for individual BMDMs along the PHATE 1 axis stimulated as indicated and calculated from data in E. (G) Scatter plots showing correlation between the PHATE 1 coordinates and the average signal intensity of all proteins secreted by each cell (left), or the number of proteins co-secreted (right) in the microwell device.

To explore the complexity in single-cell secretion responses, we visualized our high-dimensional, single-cell data in two-dimensional space using an unsupervised dimensionality reduction algorithm called PHATE (potential of heat diffusion for affinity-based transition embedding) (Moon et al., 2019). PHATE uses a diffusion mapping algorithm to visualize continual progressions, branches, and transition states within biological data. For the PHATE analysis, we included data from all four LPS dose stimulations and included measurements only from functional macrophages, which we defined as cells secreting at least one protein from Table 1. Our final PHATE analysis included secretion measurements of ten cytokines: TNF, IL-10, IL-6, IL-12p40, IL-27, CXCL1, CCL3, CCL5, GMCSF, and IFN-β. CCL2 and Chi3l3 were excluded from the analysis because they were not clearly regulated by LPS.

Projecting our single-cell data onto two dimensions revealed that most macrophages lay parallel to the horizontal PHATE 1 axis. At one end of the axis were mostly quiet cells secreting few proteins, while on the other end were macrophages secreting high levels of multiple cytokines/chemokines (C/Cs) (i.e., polyfunctional secretion; Fig. 2D). We found a strong correlation between the averaged secretion intensity across C/Cs and the PHATE 1 coordinates of individual BMDMs (Fig. 2G, left). In addition, BMDMs demonstrated a strong correlation between polyfunctionality, or the number of C/Cs being co-secreted, and their PHATE 1 coordinates (Fig. 2G, right). Taken together, the PHATE 1 axis reveals information about graded variability in the secretion response, and also captures the digital on/off response in regards to individual cytokine activation.

When we overlaid the effect of LPS dose on the placement of BMDMs along the PHATE axes, we found that unstimulated cells mostly occupied the quiet end of the PHATE 1 activation axis, characterized by low relative cytokine secretion or high secretion of only CCL3. Stimulating with 10 ng/mL LPS moved cells into the PHATE 1 activation axis, with a small population of cells secreting inflammatory C/Cs at higher intensities, though the majority of cells remained in the quiet/low activation portion. Increasing LPS dose to 100 and 1000 ng/ml led to higher numbers of cells throughout the PHATE 1 activation axis, further populating the middle and far-right end (Fig. 2E). At 1000 ng/ml LPS, the density of cells in the highest activated region of the PHATE 1 axis decreased, consistent with IL-10 negative regulation at this dose (Fig. 1B).

Estimating the kernel density of TLR4-stimulated cells across PHATE 1 identified a trimodal response with three distinguishable levels of a graded activation (inactive/low, intermediate, and high) (Fig. 2F, arrows). Examining the individual doses within the response, we observed unstimulated cells were located almost exclusively in the inactive/low activation tier, and as LPS dose increased, cells moved into the intermediate and high tiers. Notably, with the 100 ng/mL LPS stimulation dose, a greater proportion of cells resided in the high activation tier compared to the 1000 ng/mL dose. This suggests that in the presence of high levels of bacterial threat, regulatory mechanisms tailor the TLR4-stimulated dose response to avoid excessive proinflammatory activation, possibly through secretion of anti-inflammatory IL-10.

### High-dimensional analysis reveals IL-10 negative feedback modulates heterogeneity in the macrophage responder population

As mentioned above, LPS-stimulated BMDMs displayed strong correlations between their PHATE 1 coordinates and the average signal intensity across cytokines, as well as their polyfunctionality (Fig. 2G). Importantly, however, the strength of the correlation between the magnitude of secretion (i.e. signal intensity) from individual cells and the PHATE 1 coordinate varied by C/C (Fig. S2). Cytokines associated with the proinflammatory response (TNF, CCL5, CXCL1, and IL-6) exhibited the strongest PHATE 1 correlation (Pearson R ≥ 0.6), suggesting cell placement along that axis predominantly captures the degree of proinflammatory activation. In contrast, we observed weak correlation for IL-10, IL-27, and IFN-β, cytokines associated with anti-inflammatory or immuno-regulatory functions. IL-27 and IFN-β have both been implicated as paracrine signals mediating the secretion of anti-inflammatory IL-10 (Fitzgerald et al., 2013; S. S. Iyer et al., 2010). The lack of strong correlation between IL-10 and the pro-inflammatory PHATE 1 axis suggested that IL-10 may not be acting solely to resolve the pro-inflammatory response.

To explore this possibility, we next compared stimulation with LPS alone to stimulations in combination with IL-10R Ab to block IL-10 negative feedback, or in combination with recombinant IL-10 to uniformly impose IL-10 negative feedback (Fig. 3A). We overlaid the effect of stimulation dose on the placement of activated BMDMs along the PHATE axes to determine the effects of IL-10 negative feedback (Fig. 3B). Blocking IL-10 negative feedback with IL-10R Ab resulted in more highly-activated cells at the far-right end of the axis for both 100 and 1000 ng/mL in contrast to stimulating with LPS alone (Fig. 3C). However, the more significant effect of blocking IL-10 autocrine signaling was a noticeable shift in activated BMDMs from the low-responding region of the PHATE 1 axis to the more activated middle portion of the axis (Fig. 3C). In contrast, adding recombinant IL-10 dampened the proinflammatory response, with activated BMDMs exhibiting very little spread into the PHATE 1 activation axis (Fig. 3D). These data suggest that while IL-10 constrains highly activated macrophages, IL-10 negative feedback also acts to restrain low-activated cells.

**Fig. 3.**
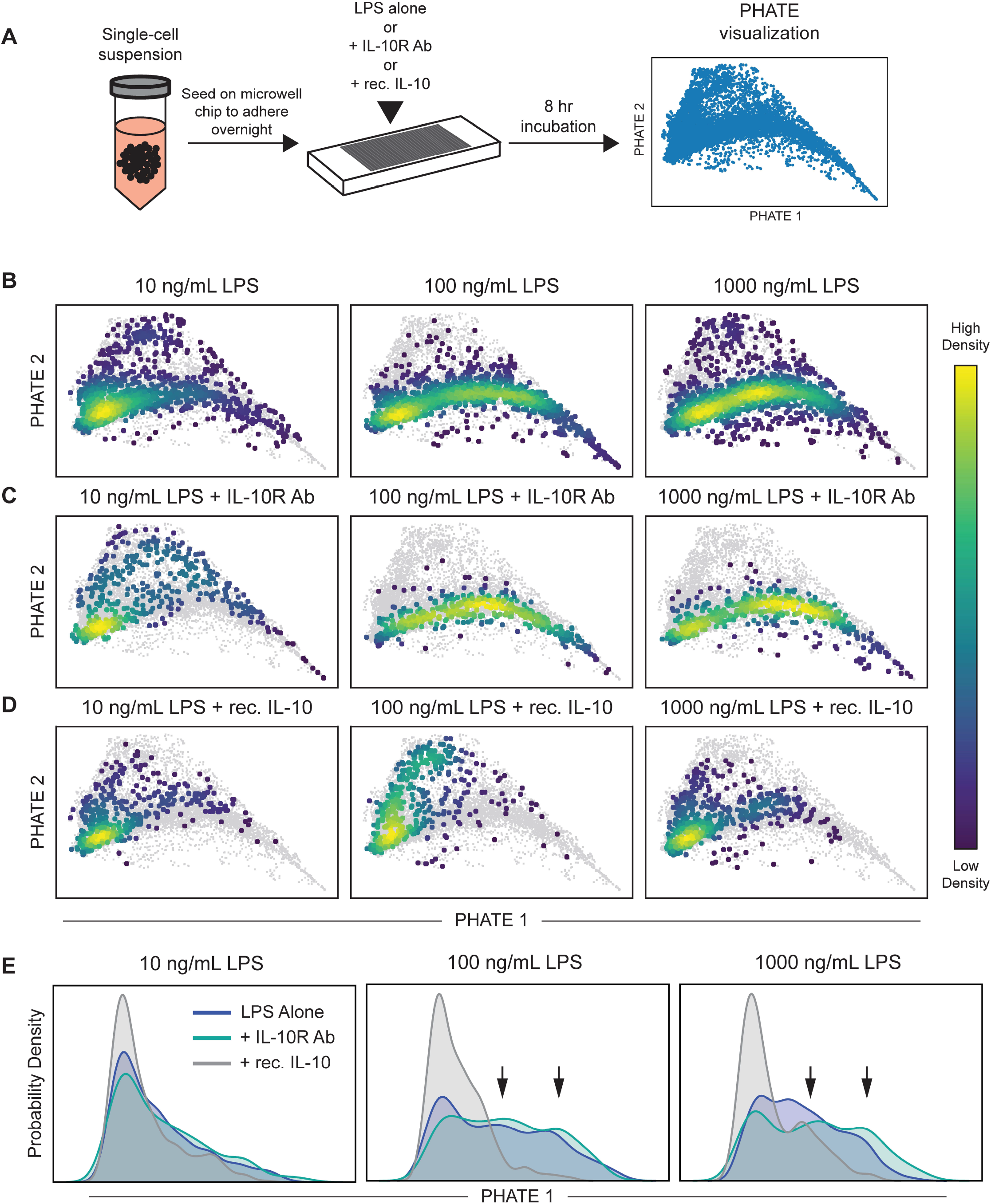
High-dimensional analysis reveals IL-10 negative feedback modulates heterogeneity in the macrophage responder population. (A) Experimental set-up for single-cell secretion profiling in the microwell device and subsequent data visualization by PHATE. (B) 2D PHATE projection of 10-dimensional single-cell secretion data of BMDMs stimulated for 8 hours with the indicated dose of LPS alone, (C) co-stimulated with LPS and IL-10R Ab (30 μg/mL), or (D) co-stimulated with recombinant IL-10 (10 ng/mL). (B-D) Data shown are colored by kernel density estimates (KDE) for the indicated subpopulation of cells (other cells greyed out for visualization). (E) Kernel density estimation for individual BMDMs along the PHATE 1 axis stimulated as indicated and calculated from data shown in B–D. Arrows indicate intermediate and high modes of the trimodal PHATE 1 KDE distribution.

To further examine this shift, we analyzed the kernel density estimate for cells along the PHATE 1 axis. We again identified the trimodal response with low, intermediate, and high levels of graded activation (Fig. 3E). Comparing cells stimulated with LPS alone to those co-stimulated with LPS and IL-10R Ab, we observed a higher proportion of cells in the low and intermediate activated states with just LPS alone. Blocking IL-10 negative feedback resulted in a decrease in the proportion of cells in the low activation region and an increase in the proportion of cells within the intermediate and high activation states. Adding recombinant IL-10 led to very high proportions of BMDMs in the low activation region at all LPS doses.

This population-wide shift in the secretion response due to perturbing IL-10 signaling was surprising, given a relatively low percentage of cells secrete high levels of IL-10 compared to other proinflammatory cytokines after LPS stimulation (Fig. 2B and C). The scope of the effect suggests IL-10 autocrine signaling can alter individual macrophage responses in the microwell device at relatively low concentrations. Notably, IL-10 perturbation had the greatest effect at the 100 ng/mL LPS dose, while the effect was much less significant at the 10 ng/mL LPS dose. This implies that IL-10 negative feedback is a dose-dependent mechanism, requiring a certain level of TLR4 activation to observe its regulation. The observation that the greatest effect of IL-10 signaling would occur at the highest levels of stimulation is consistent with our previous finding that IL-10 production is highly sensitive to stimulation dose. Overall, this analysis revealed IL-10 negative feedback regulates heterogeneity in activated BMDMs within the graded TLR4 dose response.

### Blocking IL-10 negative feedback increases TLR4 activation in low and non-responder macrophages

In our PHATE visualizations, we observed the greatest impact of IL-10 autocrine signaling on cells in the low-responding region of the activation axis. Given this result, we speculated that low and nonresponding cells may be primary targets of IL-10 negative feedback. This effect could be direct, and could also act indirectly via IL-10 negative regulation of TNF activation, which positively regulates other cytokines (Fig. 1A) (Caldwell et al., 2014; Xue et al., 2015).

To explore the effect of IL-10 on TNF, we converted single-cell secretion intensities from the microwell device to individual protein concentration values using recombinant protein standard curves (Fig. S4). Concentrations below the threshold of detection in the microwell device were set to 1 pg/ml (i.e., 0 on a log scale) for visualization, which was just below the lowest extrapolated concentrations detected in individual cells. Probability histograms showing single-cell TNF secretion in the microwell device revealed that blocking IL-10 signaling increased the percentage of cells secreting TNF, and therefore conversely decreased the non-responder population (Fig 4A). The IL-10 mediated increase in the activated population is a result of non-activated macrophages becoming low-level TNF secretors, rather than activated cells secreting more TNF. In contrast, adding recombinant IL-10 significantly decreased the percentage of cells responding to LPS stimulation and the average level of secretion amongst responding cells, and thus greatly increased the TNF non-responder subpopulation.

**Fig. 4.**
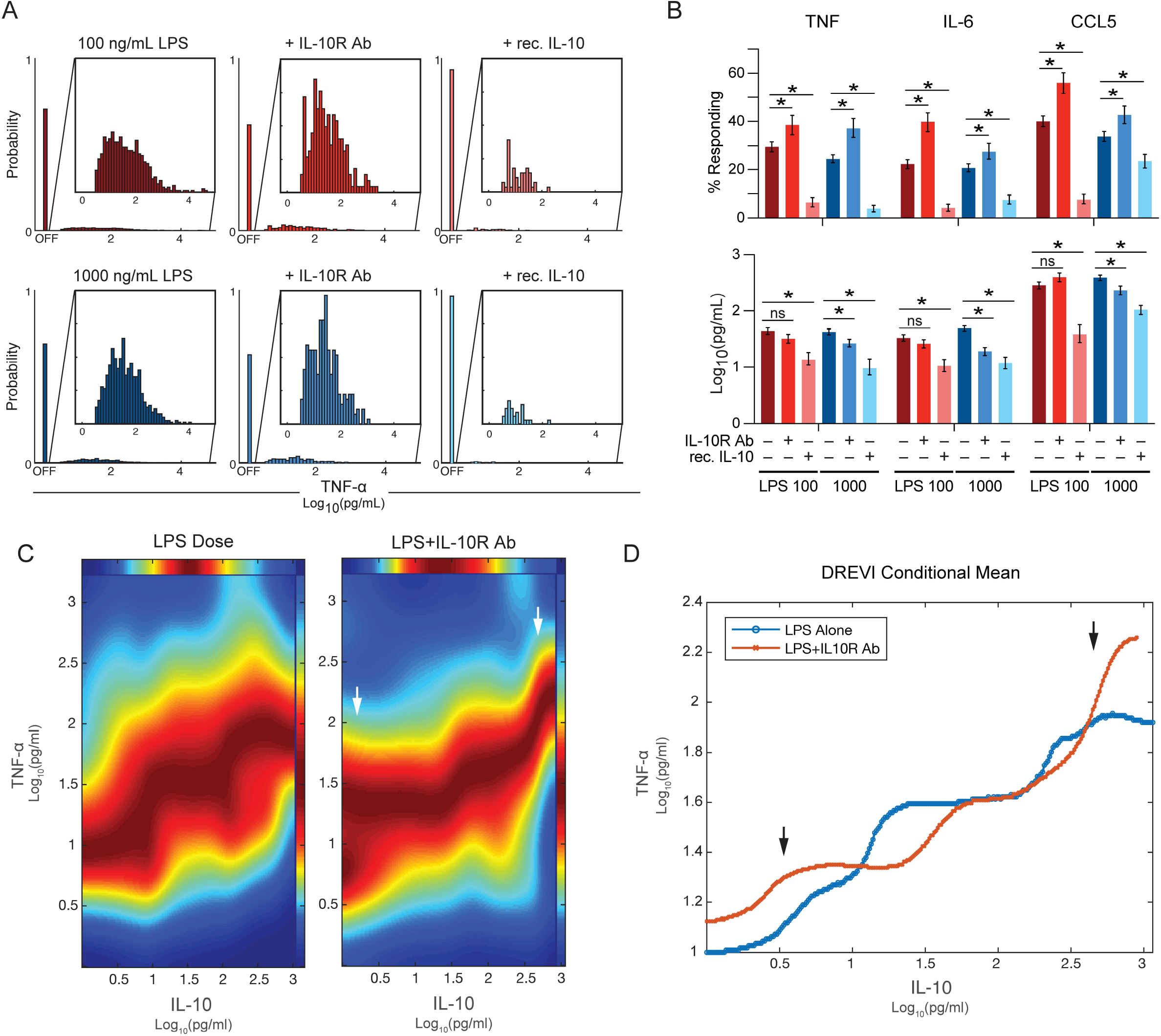
Conditional-density visualization reveals low and high levels of IL-10 negative feedback differentially modulates proinflammatory activation in TLR4-stimulated macrophages. (A-D) Fluorescent intensities from the microwell assay were converted to protein concentrations using recombinant protein standard curves (See materials and methods for details). LPS alone data are pooled from 3 independent experiments, IL-10R Ab data pooled from 2 independent experiments, rec. IL-10 data are one representative experiment. (A) Probability histograms for TNF secretion after 8 hours of the indicated stimulation in the microwell device. Non-responding cells were set to 1 for visualization (labeled “OFF”). Inset graphs show only the responding (TNF+) subpopulation. (B) Bar graphs depict the average percentage of cells secreting the indicated cytokine/chemokine above threshold in response to the indicated stimulation cues after 8 hours in the microwell assay (top), or mean secretion level from responding subpopulation of cells (only cells secreting indicated cytokine/chemokine above detection threshold in microwell device) (bottom). Data shown are mean with 95% confidence intervals calculated by bootstrapping. Significance determined by non-overlap of confidence intervals. (C and D) Conditional density re-scaled visualization (DREVI) plots showing the relationship between IL-10 and TNF. Data shown are pooled to include all stimulation doses (0, 10, 100, 1000 ng/mL LPS with or without IL-10R Ab at 30 μg/mL) where the 2 indicated cytokines were co-secreted during the 8-hour incubation (LPS alone: n=516, +IL-10R Ab: n=212). Arrows indicate areas of low and high IL-10 secretion most affected by IL-10 negative feedback. (D) Edge response functions for DREVI plots shown in C were fit according to the conditional mean at the region of highest conditional density.

We further examined the protein concentration data for IL-6 and CCL5, and calculated the percentage of cells secreting each individual cytokine above threshold, as well as the average protein secretion across the responding cell population (excluding non-responding cells). Similar to its effect on TNF, blocking IL-10 negative feedback significantly increased the percentage of cells producing IL-6 and CCL5, while in contrast, adding recombinant IL-10 greatly reduced the percentage of cells secreting these proinflammatory cytokines (Fig. 4B). However, we found blocking IL-10 negative feedback had no effect on the average level of secretion in the responder population at 100 ng/mL LPS, and even decreased the average level of secretion in the responder population at 1000 ng/mL LPS stimulation (Fig. 4B), presumably due to recruitment of low-level secretors into the activated population. These data suggest that IL-10 negative feedback acts to block the total number of cells secreting TNF and other proinflammatory cytokines, rather than acting solely to reduce proinflammatory secretion in activated cells.

### Conditional-density visualization reveals low and high levels of IL-10 negative feedback differentially modulate proinflammatory activation in the TLR4 response

A surprising observation upon blocking IL-10 negative feedback is that the affected responder population is larger than the subpopulation of cells secreting IL-10. For example, at 100 ng/mL LPS stimulation, an average of 13.1% [11.6, 14.7] (95% CI) of cells secreted IL-10 above background in the microwell device (Fig. 2C). However, blocking IL-10 negative feedback at this same dose caused the percentage of cells secreting IL-6 to increase 17.6%[15.1, 19.8], and the percentage of cells secreting CCL5 to increase 16.1% [14.4, 17.8] (Fig. 4B). This discrepancy suggests IL-10 may elicit negative feedback at concentrations below the limit of detection in the microwell device.

To investigate this further, we examined how the concentration of secreted TNF varied with the concentration of secreted IL-10 using conditional density re-scaled visualization (DREVI) (Krishnaswamy et al., 2014). DREVI rescales multiplexed single-cell data points by their conditional density in order to robustly visualize relationships between proteins across a larger dynamic range. In this case, DREVI allowed us to emphasize the small number of cells that exhibited low levels of IL-10 secretion (i.e., < 10 pg/ml) and look at the effect of this secretion on TNF. We pooled our interpolated microwell secretion data from all stimulation doses (0, 10, 100, 1000 ng/mL LPS) with and without IL-10 receptor blocking, and then we confined our DREVI analysis to individual BMDMs producing both cytokines.

DREVI analysis of the co-secretion relationship between TNF and IL-10 after LPS stimulation identified three tiers of activation (low, intermediate, and high), similar to PHATE. Within each tier, as IL-10 increased, the concentration of TNF plateaued or decreased (Fig. 4C, left). Blocking IL-10 negative feedback altered the functional relationship between these two cytokines, specifically at the lowest concentrations of IL-10 (< 10 pg/ml) and highest concentrations of IL-10 (> 300 pg/ml; Fig. 4C, right). With LPS alone, we observed low levels of TNF were co-produced with low levels of IL-10; similarly, intermediate levels of TNF were co-produced with intermediate levels of IL-10, and high levels of TNF with high levels of IL-10. In comparison, without IL-10 negative feedback, we observed co-production of low levels of IL-10 secretion with both low and intermediate levels of TNF secretion. This suggests that low concentrations of IL-10 negative feedback prevented low TNF-responders (or non-responders) from becoming intermediate responders. Further, without IL-10 negative feedback we observed the highest tier of TNF activation no longer plateaued with increased IL-10, but instead continued to increase (Fig. 4D).

Examining the co-secretion relationship between IL-6 and IL-10 in BMDMs stimulated with LPS alone revealed only two levels of IL-6 activation: intermediate and high (Fig. S5A, left). We observed low concentrations of IL-10 co-produced with intermediate levels of IL-6, and then a sharp increase in IL-6 secretion coinciding with intermediate IL-10 secretion (> 50 pg/ml). At the highest concentrations of IL-10 secretion, we saw the levels of co-produced IL-6 dramatically decrease back down to intermediate levels, consistent with IL-10-mediated negative feedback. In contrast, without IL-10 receptor signaling, we observed a fraction of cells co-producing high levels of IL-10 and high levels of IL-6 (Fig. S5A, right and Fig. S5B). Interestingly, we still observed a fraction of cells with decreased IL-6 secretion at high IL-10 concentrations, even when IL-10 signaling was blocked, suggesting there are IL-10-independent negative feedback mechanisms that also act to resolve high IL-6 responders following TLR4 activation.

In addition, blocking IL-10 signaling resulted in a new population of cells exhibiting low IL-6 activation at low concentrations of IL-10 (Fig. S3A, right; Fig. S3B). As mentioned above, the DREVI plots only include cells secreting both IL-6 and IL-10, which means non-responding cells are not visualized in the DREVI plots. Given the observation that blocking IL-10 negative feedback almost doubles the percentage of cells that secrete IL-6 (Fig. 4B), it is likely that additional cells that were non-responders (and thus not shown) in the DREVI plot for LPS alone, are now included and create a new tier of cells secreting low concentrations of IL-6. Overall, our data establishes that low concentrations of IL-10 are sufficient to regulate heterogeneity in the TLR4 response.

### IFN-β largely mediates IL-10 regulation of the threshold for TLR4 activation, while TNF positive feedback activates the resolving role of IL-10

IL-10 is induced by LPS and also by paracrine signaling from other LPS-induced cytokines in macrophages, specifically TNF and IFN-β (Fig. 1A). To determine to what extent the IL-10 negative feedback regulation we observed is mediated by paracrine signaling of TNF and IFN-β, we again performed multiplexed single-cell secretion profiling on BMDMs stimulated with 100 ng/mL LPS alone, or costimulated with soluble TNF receptor (sTNFR) to block TNF autocrine signaling, or an interferon-α/β receptor blocking antibody (IFNAR Ab) to block IFN-β autocrine signaling. We confirmed that both sTNFR and IFNAR Ab significantly reduced the fraction of cells secreting IL-10 as well as the magnitude of that secretion (Fig. 5C and S6), confirming roles for TNF and IFN-β in promoting IL-10 production.

**Fig. 5.**
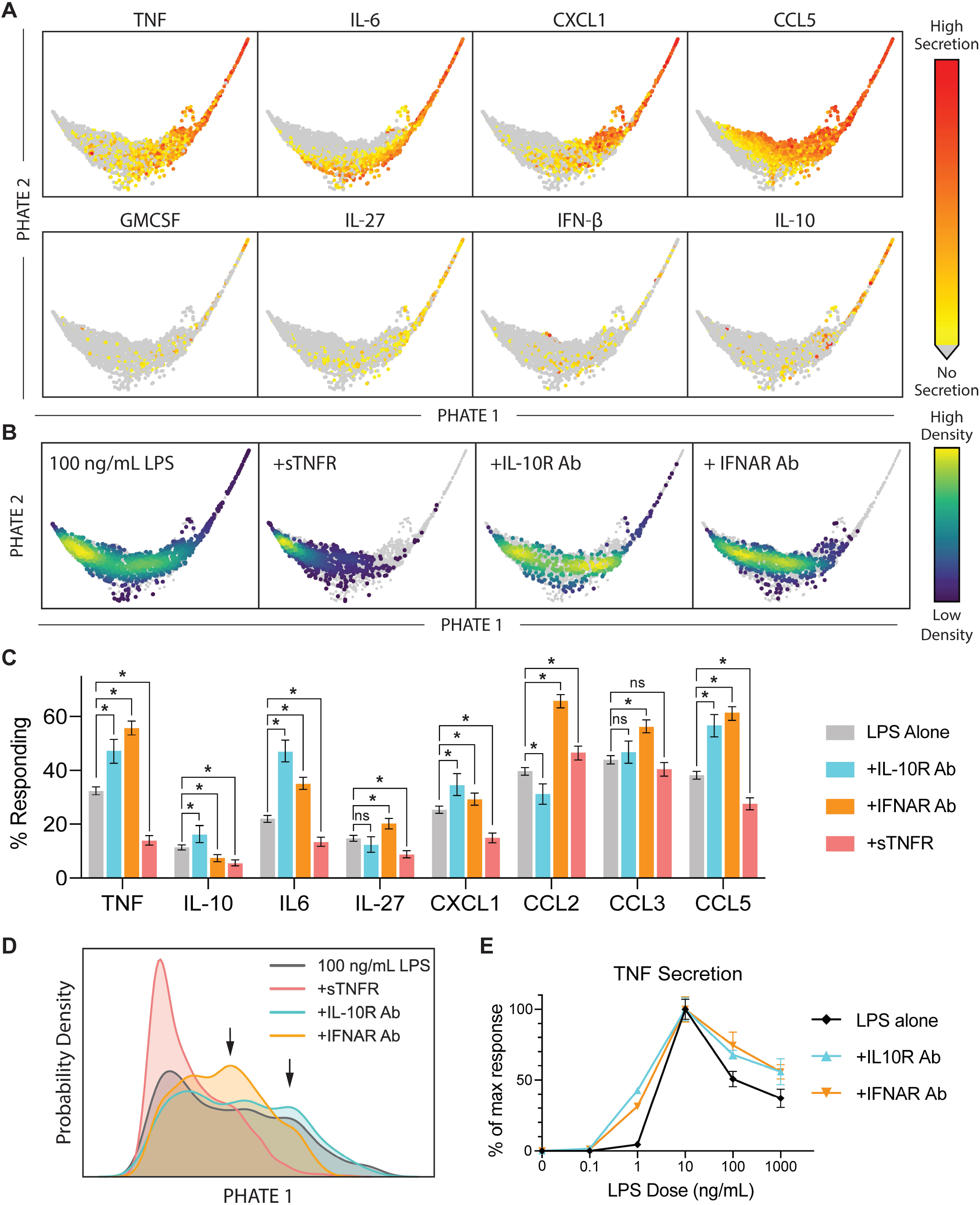
IFN-β largely mediates IL-10 regulation of the threshold for TLR4 activation, while TNF positive feedback activates the resolving role of IL-10. 2D PHATE projection of 10-dimensional singlecell secretion data from BMDMs stimulated with 100 ng/mL LPS or co-stimulated with IL-10R Ab (30 μg/mL), IFNAR Ab (5 μg/mL), or sTNFR (5 μg/mL) for 8 hours in the microwell device. (A) Data are colored by relative secretion intensity of the indicated protein. (B) Data shown are colored by cell density for the indicated subpopulation of cells (other cells greyed out for visualization). (C) Bar graphs show percentage of cells secreting the indicated cytokine/chemokine above threshold in response to the indicated stimulation cues after 8 hours in the microwell assay. Data shown are mean with 95% confidence intervals calculated by bootstrapping. Significance determined by non-overlap of confidence intervals. (D) Kernel density estimation for individual BMDMs along the PHATE 1 axis stimulated as indicated and calculated from data shown in A–B. Arrows indicate intermediate and high modes of the trimodal PHATE 1 distribution. LPS alone data pooled from 3 independent experiments. IL-10R and IFNAR data are each one representative experiment. sTNFR data pooled from 2 independent experiments. (E) BMDMs were stimulated with the indicated doses of LPS alone or co-stimulated as indicated for 24 hours, after which cell supernatants were collected and TNF production measured by ELISA. Data shown are mean ± SEM of 2 biological replicates.

We combined our sTNFR and IFNAR Ab results with our previous IL-10R Ab experiment, and visualized the high-dimensional secretion data with PHATE. Co-stimulation with sTNFR substantially diminished the proinflammatory response, as evidenced by the lack of cells throughout the PHATE 1 proinflammatory axis (Fig. 5A-B), and by significant decreases in the percentage of cells producing inflammatory cytokines IL-6, IL-27, CXCL1, and CCL5 (Fig. 5C). These results are consistent with the role of TNF as a strong positive regulator of the TLR4 response. This loss of this positive feedback prevented a direct analysis of the effect of IL-10 reduction on the activation of TLR4-stimulated responder macrophages.

In contrast, cells co-simulated with IFNAR increased their proinflammatory activity, as evidenced by movement of cells farther into the proinflammatory activation axis (Fig. 5B), and by significant increases in the percentage of cells secreting TNF, IL-6, CXCL1, and CCL5, which was similar to what we observed when blocking IL-10 signaling via co-stimulation with anti-IL10R Ab (Fig. 5C). There were, however, differences in how blocking IL-10 signaling versus IFN-a/β signaling affected secretion of individual proteins. For example, the fraction of BMDMs secreting IL-27 and CCL3 remained unchanged with IL-10 signaling blocked, but significantly increased when IFNAR was blocked. Despite these differences, overall, we found significant functional crossover between negative feedback from IL-10 and IFN-β in both the digital and analog components of the TLR4 response.

We again analyzed the kernel density estimate for cells along the PHATE 1 axis to further visualize the effects of blocking different paracrine signals. Blocking TNF moved most BMDMs into the lowest tier of TLR4 activation, consistent with its primary role in positive feedback. IFNAR blocking resulted in a significant increase in BMDMs mostly in the intermediate tier of activation, but did not increase activation in the highest tier. In contrast, blocking IL-10R increased BMDMs in both the intermediate and high activation tiers (Fig. 5D). Taken together, these data raise the possibility that TNF, or another paracrine signal, is promoting IL-10 secretion primarily in the highest activation tier.

Our evidence that IL-10 and IFN-β negative feedback largely act to restrain cells from responding to LPS led us to postulate that IL-10 may play a role in setting the threshold for TLR4 activation. A dosedependent threshold for TLR4 activation has been previously described (Gottschalk et al., 2016) and was clearly evident in the activation of TNF secretion (Fig. 1C and S1). To explore a role for IL-10 in setting this threshold, we stimulated BMDMs in a plate with increasing doses of LPS alone or co-stimulated with IL-10R Ab or IFNAR Ab. After 24 hours, we collected cell supernatants and measured TNF concentration by ELISA. We found that with LPS stimulation alone, we observed TNF induction above 1 ng/mL LPS stimulation dose, but with IL-10 or IFN-β extracellular signaling blocked, we saw TNF induced between 0.1 and 1 ng/mL LPS (Fig. 5E). This data show that IL-10, mediated by IFN-β paracrine signaling, contributes to a threshold of TLR4-induced proinflammatory activation.

## Discussion

Macrophages exhibit heterogeneous inflammatory secretion (Lu et al., 2015; McWhorter et al., 2016), while simultaneously responding to the cytokines they secrete and the cytokines secreted by neighboring cells (Gautier et al., 2005; Rand et al., 2012; Xue et al., 2015). To ensure a response that is targeted for the appropriate level of immune threat while avoiding tissue damage due to hyperinflammation, macrophages use paracrine communication networks (Peter J. Murray & Smale, 2012). The secretion of anti-inflammatory IL-10 following TLR4 stimulation is a negative feedback mechanism that combats hyperinflammation. In this study, we used a combination of single-cell and population assays to investigate the role of IL-10 negative feedback in implementing the heterogeneous TLR4 secretion response.

First, our results confirmed that the potent immunosuppressive effect of IL-10 secretion in TLR4-stimulated macrophages is both time and dose-sensitive (Fig. 1B and 1C). This time sensitivity is supported by literature that shows production of paracrine signals such as IFN-β and IL-27 are required for robust TLR4-induced IL-10 production, resulting in a time delay (E. Y. Chang et al., 2007; Ernst et al., 2019; S. S. Iyer et al., 2010). Further, the switch-like dose response we observed in IL-10 and TNF secretion induction at both 8 and 24 hours provides evidence that a previously identified switch-like activation threshold in upstream MAPK signaling is further perpetuated at the level of functional output (Fig. 1C and S1). While these thresholds exemplify a cell-intrinsic mechanism of dose discrimination, the inverse relationship between TNF and IL-10 at high LPS doses indicates proinflammatory activation is further influenced by extracellular signals and cell-cell communication.

High-dimensional analysis of single-cell secretion data revealed a graded response along an extended axis of proinflammatory secretion encompassing both the digital (on/off) and analog (graded) activation components (Fig. 2). Using the dimensionality reduction algorithm PHATE (Moon et al., 2018) we observed correlation between single-cell coordinates along the horizontal PHATE 1 axis and secretion intensity of several proinflammatory cytokines (Fig. S2). Distinctly, secretion of IL-10, IL-27, and IFN-β did not correlate strongly with the PHATE 1 axis (Fig. S2), suggesting that regulation of immunomodulatory cytokines is separate from proinflammatory cytokines, and that these regulatory decisions vary across individual cells. This diversification is likely another strategy used by macrophages to restrain proinflammatory secretion.

Using kernel density estimation along the PHATE 1 axis, we identified trimodal activation of proinflammatory secretion, a pattern that was even clearer after blocking IL-10 negative feedback (Fig. 2 and Fig. 3). Previous studies have identified bimodal cytokine production when measuring production of a single protein, such as TNF or IL-6 (Muldoon et al., 2020; Shalek et al., 2013), which is consistent with our observed intermediate and high activation tiers. Our analysis expands on those observations by simultaneously measuring 12 cytokines and chemokines that are co-produced in the TLR4 response. This multiplexed analysis revealed an additional mode of low activation, in which cells secrete few cytokines at low intensity, or only robustly secrete chemokines such as CCL3.

Previous work in our lab identified functional clusters within TLR-stimulated macrophages instead of activation tiers within a continuum. Specifically, our study of U937 macrophages and human MDMs identified loosely defined functional subgroups after stimulation with various TLR agonists (Lu et al., 2015). Notably, the majority of these subgroups were defined based on varied degrees of activation as opposed to the secretion of entirely different proteins. It follows that our inclusion of additional doses of TLR4 agonist instead of a single dose allowed further visualization of the full range of macrophage activation, and we hypothesize some of the previously identified subgroups may have represented snapshots of different activation tiers as opposed to functionally distinct clusters.

Our study further shows how the TLR-induced macrophage secretion response is regulated through both positive and negative feedback motifs mediated through paracrine signals. Multiplexed single-cell secretion profiling revealed how blocking IL-10 signaling with IL-10R Ab primarily suppressed activation in low and non-responding cells, with a minimal effect on the highest activated cells (Fig. 3). This widespread effect of IL-10 was disproportionate to the level of IL-10 secretion we observed, and using DREVI, a novel analysis method that specializes in identifying relationships between proteins that may be obscured by insufficient sampling density (Krishnaswamy et al., 2014), we confirmed low concentrations of IL-10 were responsible for negative feedback in the low-activation macrophage population (Fig. 4 C and D). Similarly, DREVI confirmed high levels of IL-10 were responsible for tamping down inflammatory cytokine production in the high activation macrophage population. These results confirm a dual role for IL-10 in both restraining and resolving the TLR4-induced macrophage secretion response.

While IL-10 acts via paracrine signaling to negatively regulate the TLR4 response, there is an additional layer of regulation in that IL-10 secretion is induced by the paracrine signals TNF and IFN-β. Our results expand on the complexity of this regulation, revealing that TNF and IFN-β appear to mediate different functional roles of IL-10 within the TLR4-stimulated macrophage response (Fig. 5). Blocking IFN-β signaling specifically increased the fraction of macrophages in the intermediate activation tier but did not affect macrophages in the high activation tier, in contrast to blocking IL-10 signaling, which affected both (Fig. 5D). This suggests that IFN-β-mediated IL-10 restrains cells at the low end of the proinflammatory axis, while other paracrine signals-TNF and possibly others–mediate IL-10 negative regulation of cells in the highest activation tier (Fig. 6).

**Fig. 6.**
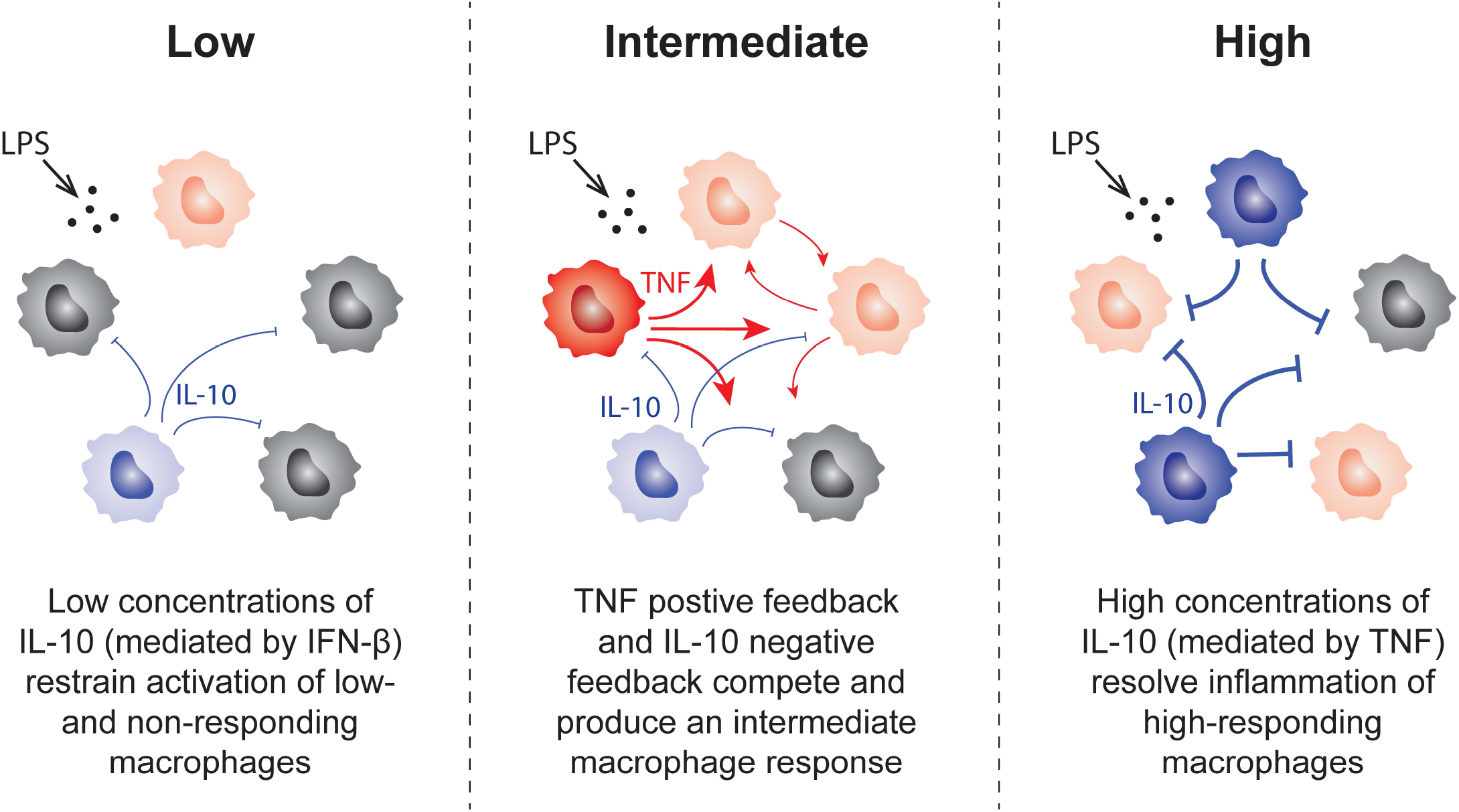
IL-10 plays a dual role in restraining and resolving the TLR4-induced inflammatory response. Schematic model illustrating dual roles for IL-10 in the TLR4 response. At low levels of TLR4-induced activation, low concentrations of IL-10 secretion mediated by intermediate IFN-β signaling restrain proinflammatory activation of low and non-responding macrophages by raising the activation threshold (left). At high levels of TLR4-induced proinflammatory activation, high concentrations of IL-10 negative feedback mediated by intermediate TNF signaling act to resolve highly-active proinflammatory cells and avoid overresponse (right).

Building on our observations from the single-cell data, we demonstrated that IL-10 and IFN-β elicit their negative feedback by raising the threshold for TLR4 activation in a population of macrophages (Fig 5E). While the relationship between IL-10 and IFN-β and their anti-inflammatory function have been well documented in literature (E. Y. Chang et al., 2007; McNab et al., 2014; Shalek et al., 2014), a recent study of BMDMs stimulated with varied classes of bacteria demonstrated IFN-β negative feedback mechanisms independent of IL-10 (Gottschalk et al., 2019). Our analysis was unable to completely delineate IFN-β negative feedback from IL-10 negative feedback; therefore, further studies are necessary to illuminate their distinct roles in the anti-inflammatory response.

In contrast to IFN-β, reducing TNF paracrine signaling with sTNFR resulted in a significant reduction in the secretion of proinflammatory cytokines from individual TLR4-stimulated macrophages in the microwell device (Fig. 5). Blocking TNF signaling limited the number of macrophages that entered the intermediate activation tier, and prevented macrophages from reaching the high activation tier. This finding builds upon ample research on TNF positive feedback enhancing secretion of other proinflammatory cytokines (Covert, 2005; Werner, 2005; Xue et al., 2015). Further, our results show the loss of TNF positive feedback via autocrine signaling has a larger effect on the overall response than the reduction in autocrine IL-10 negative feedback. Interestingly, previous work in our lab found co-stimulating macrophages with LPS and recombinant TNF to create a uniform TNF signal did not amplify proinflammatory secretion (Xue et al., 2015). This suggests there is a limit to proinflammatory amplification from TNF positive feedback, and the interaction of positive and negative feedback motifs enables fine-tuning of the overall inflammatory response.

Our finding that TNF is a major autocrine regulator of the TLR4 response is in contrast to Caldwell et al., which reported that in response to LPS, TNF engaged exclusively in paracrine signaling to neighboring, non-macrophage cells (Caldwell et al., 2014). The authors argued that weaker TLR stimuli such as CpG (TLR9 agonist) exhibited TNF-enhanced NF-κB activity, but the speed and duration of TLR4-stimulated NF-κB activity eliminated the possibility for autocrine enhancement. One reason for the differences in our studies could be due to the loss of paracrine signaling in the microwell device. As previously described, paracrine signaling amplifies macrophage TLR4 responses (Xue et al., 2015), and therefore in the microwell device the TLR4 response may be weaker. It follows that the effect of sTNFR may then be amplified due to this lack of additional TNF signals from neighboring cells. We speculate that the weaker response in the microwell device induces levels of NF-κB activation that can be amplified by TNF autocrine signaling.

This work highlights the importance of combining both population and single-cell analysis methods to form a more complete picture of immune cell behavior. While great progress has been made in singlecell analysis methods, further work must be done to improve and expand the options for single-cell functional secretion analysis. There are limits to the regulatory mechanisms that can be uncovered in our microwell device. For instance, our device cannot capture dynamics of macrophage secretion. Our assay measures cumulative secretion over time, and it is possible that proteins that appear to be co-secreted are in reality sequentially secreted. Parsing the specifics of that polyfunctionality would require further assay development. Additionally, the microwell device cannot investigate spatial regulation within the TLR4 response. Cell density and quorum sensing have been cited as additional regulators of macrophage activation (Muldoon et al., 2020), and further illumination of these mechanisms would inform higher order immune regulation and deepen our understanding of macrophage behavior in vivo.

Overall, our research revealed important aspects of the sophisticated network guiding population-level macrophage secretion responses, and furthered our understanding of IL-10 and the interplay between positive and negative feedback motifs (Fig. 6). This analysis informs higher order immune regulation and could pave the way for improved cytokine immunotherapies for autoimmune diseases.

## Acknowledgments

We thank Andre Levchenko, Smita Krishnaswamy, and David Hafler, as well as members of the Miller-Jensen lab for insightful advice and helpful discussion. We also thank I. Kelsey for critical reading of the manuscript. This work was supported by the National Institutes of Health (R01-GM123011 to K.M.J., and RO1-GM072024, to A.L.), and the National Science Foundation (DGE1122492).

## Author Contributions

Conception: A.F. Alexander, and K. Miller-Jensen. Development of methodology: A.F. Alexander and K. Miller-Jensen. Acquisition of data: A.F. Alexander, H. Forbes. Analysis and interpretation of data: A.F. Alexander and K. Miller-Jensen. Writing, review, and/or revision of the manuscript: A.F. Alexander and K. Miller-Jensen. Study supervision: K. Miller-Jensen.

## Materials and Methods

### Mouse breeding and cell culture

All mice were housed in the Yale Animal Resources Center in specific pathogen-free conditions. All animal experiments were performed according to the approved protocols of the Yale University Institutional Animal Care and Use Committee. Wild type C57BL/6J mice were purchased from Jackson Laboratories. Bone marrow derived macrophages were generated as previously described (Trouplin et al., 2013). In brief, bone marrow was extracted from the tibias and femurs of the hind legs of mice with a syringe. Afterwards, red blood cells were lysed with ammonium-chloride-potassium lysis buffer (Lonza) and the cells were incubated for 4 hours at 37°C with 5% CO2 in a non-tissue culture treated petri-dish with BMDM media (RPMI supplemented with 10% FBS, 100 U/mL penicillin, 100 μg/mL streptomycin, 1% sodium pyruvate, 25 mM HEPES buffer, 2mM L-glutamine, and 50 mM 2-mercaptoethanol). After 4 hours, the non-adherent cells were transferred to a new non-tissue culture treated petri dish and incubated with BMDM media supplemented with 20 ng/mL M-CSF (PeproTech). On day 3 after plating, 10 mL of BMDM media supplemented with 20 ng/mL M-CSF was added to the dish. After 6 days, BMDMs were lifted with PBS+5mM EDTA and gentle scraping. Cell suspensions were seeded onto plates at a density of 250,000 cells/ml for population ELISA experiments or onto the microwell device at a density of 125,000 cells/mL.

### In vitro treatments and quantification of secretion in population

BMDMs were plated and stimulated in BMDM media with 10 ng/mL M-CSF (PeproTech). Cells were plated at the appropriate density (described above) and allowed to adhere overnight. Cells were stimulated with LPS (Invivogen) alone, or co-stimulated with LPS and IL-10R blocking antibody (BioLegend, clone: 1B1.3a), or LPS and recombinant IL-10 (PeproTech), or LPS and IFNAR1 blocking antibody (Invitrogen, clone: MAR1-5A3) for the times indicated in the legends. Cell culture medium was collected at the end of the incubation period and was assayed by ELISA according to the manufacturer’s instructions. The antibody pairs used in the ELISAs were the same as those used for the microwell assay and can be found in Table 1.

### Microwell assay for single-cell secretion profiling

Single-cell secretion profiling experiments were performed as previously described (Lu et al., 2013; Xue et al., 2015) (Appendix A). Briefly, capture antibodies (Table 3.1) were flow patterned onto epoxy silane-coated glass slides (Super-Chip; ThermoFisher). The polydimethylsiloxane (PDMS) microwell arrays and antibody barcode glass slides were blocked using complete RPMI. BMDMs were suspended in complete RPMI supplemented with 10 ng/mL M-CSF and were added to the PDMS microwell array and allowed to adhere overnight. The next day BMDMs were stimulated as indicated with complete RPMI supplemented with 125 nM of live cell marker (Calcein AM; ThermoFisher) to allow automatic live cell detection. The BMDMs in the PDMS microwell array were then covered with the antibody barcode slide, secured with plates and screws, and allowed to incubate for 8 hours. At the end of the incubation period, the device was imaged with an automated inverted microscope (Axio Observer; ZEISS) to record well position and cell locations. The device was then disassembled and the sandwich immunoassay was performed: The glass slide was incubated with a mixture of detection antibodies (Table 3.1) for 2 hours, followed by incubation with 20 μg/mL streptavidin-APC (eBioscience) for 30 minutes, rinsed with PBS and deionized water, and scanned with a Genepix 4200A scanner (Molecular Devices).

### Single-cell secretion profiling and data processing

Device images were analyzed using a custom script in MATLAB (MathWorks) to automatically detect well location and number of cells per well, extract all signals from each well, and process the data (https://github.com/Miller-JensenLab/Single-Cell-Analysis). In brief, after automatic well and live cell detection, signal image registration, and manual curation, the software automatically extracted the intensity signal from each antibody for all the microwells in the device. This signal across the chip for each individual antibody was normalized by subtracting a moving Gaussian curve fitted to the local zero-cell well intensity values. A secretion threshold for each antibody was set at the 99th percentile of all normalized zero-cell wells. Data was transformed using the inverse hyperbolic sine with cofactor set at 0.8x secretion threshold.

### Multidimensional PHATE analysis

To further visualize high-dimensional secretion data from the microwell assay, the dimensionality reduction algorithm known as potential of heat diffusion for affinity-based transition embedding (PHATE) was used. PHATE analysis was performed using the available MATLAB and Python packages from the Krishnaswamy lab (https://github.com/KrishnaswamyLab/PHATE). Standard parameters were used to make 2-dimensional PHATE projections from the 10-dimensional BMDM dataset. Manual adjustments to the t parameter were made for optimal visualization. PHATE plots were colored by density or by relative secretion levels of each protein as indicated; cells below the secretion threshold were greyed out for visualization. Extracted PHATE parameters were analyzed using custom software written in MATLAB and Python.

### Recombinant protein standard curves

To convert measured fluorescent intensities from the microwell assay to concentrations of cytokines and chemokines, we used recombinant protein calibration curves (Fig. S3.4). Recombinant protein standard curves were derived by measuring the intensity values of recombinant proteins of concentrations between 39 and 5000 pg/mL in the microwell assay. The 4 Parameter Logistic (4PL) nonlinear regression model was used to fit the standard curves, and the 95% confidence intervals were calculated with Prism8 (GraphPad). Concentration values for intensities less than the detection limit of the calibration curve were set to zero or set to 1 for histogram visualization. For further analysis, interpolated concentration data from the microwell assay was analyzed using conditional density re-scaled visualization (DREVI). The DREVI software was used to visualize 2D stochastic relationships in single-cell data using conditional density estimation, representing how a variable X influences a variable Y, by depicting the distribution of Y for each value of X. DREVI algorithm was downloaded from the Pe’er Lab website (https://dpeerlab.github.io/dpeerlab-website/dremi.html).

### Statistical analysis

Data were presented as means ± SEM unless otherwise specified. Statistical analysis was performed by ordinary 2-way ANOVA and the Dunnett method for correction of multiple comparisons as specified in the figure legends. All analyses were performed using Prism 8.4.1 software (GraphPad). For single-cell distributions, statistics were performed using a bootstrapping procedure to calculate the confidence intervals associated with sampling error in single-cell data. To obtain confidence intervals through bootstrapping, the single-cell datasets for each condition were sampled 10,000 times with replacement, and the metric of interest was calculated for each resampled dataset. We then calculated a 95% confidence interval for these resampled datasets, and statistical significance was assigned to pairwise comparisons with non-overlapping confidence intervals. This bootstrapping procedure was done using a custom script in MATLAB.

