## Supplemental Figures for "Single-cell secretion analysis reveals a dual role for IL-10 in restraining and resolving the TLR4-induced inflammatory response"

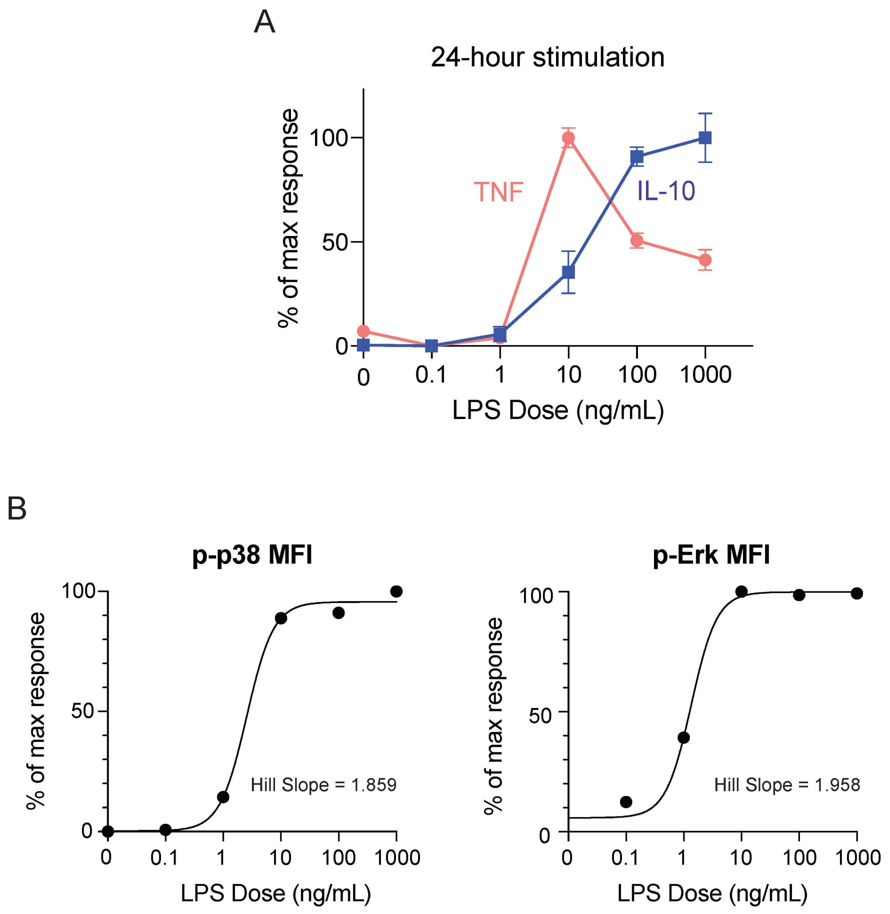


**Figure S1. (related to Fig. 1) TLR4-induced proinflammatory activation in macrophages is dose dependent.** (A) BMDMs were stimulated with the indicated doses of LPS for 24 hours, after which cell supernatants were collected and TNF and IL-10 concentration measured by ELISA. (B) BMDMs were stimulated with the indicated concentration of LPS for 15 minutes, after which MAPK activation via p-p38 (left) and p-ERK (right) were measured by flow cytometry. 4-parameter logistic (4PL) dose-response curves were fit to the percent of maximum median fluorescent intensity (MFI). Data include one representative experiment for each target.

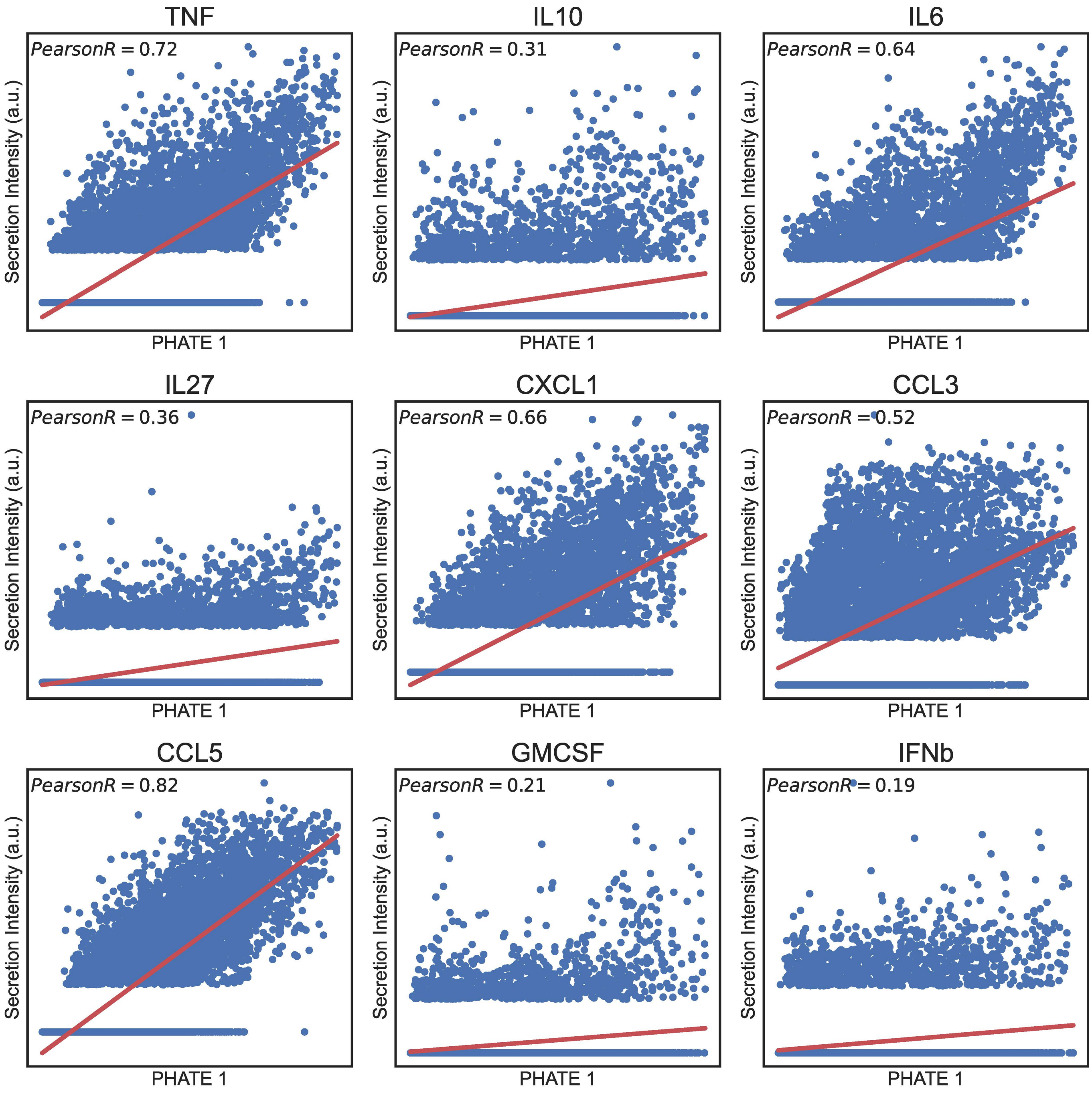


**Figure S2 (related to Fig. 2) Cytokine secretion intensity in the microwell device correlates with the PHATE 1 axis.** Scatter plots show correlation between single-cell secretion intensity for the indicated cytokines from BMDMs stimulated as described in Fig. 2, and the PHATE 1 coordinates extracted from 2D PHATE embeddings of the data. Best fit line shown in red.

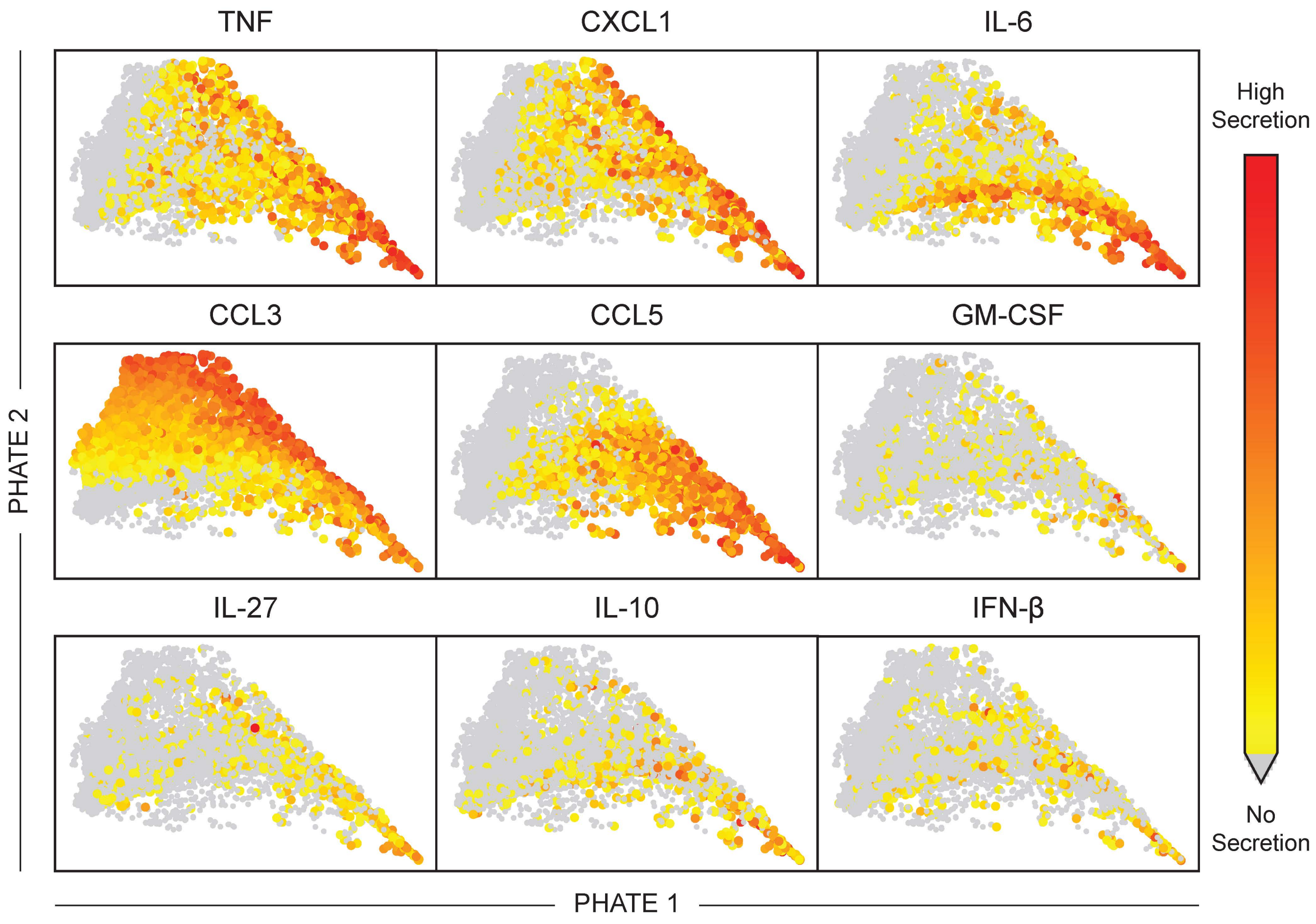


**Figure S3 (related to Fig. 3) TLR4-activated macrophages lie along a single, graded proinflammatory axis.** 2D PHATE visualization of single BMDMs based on the secretion levels of 10 proteins after stimulation as described in Fig. 3. Data are colored by relative secretion intensity of indicated protein, arbitrary units (a.u.)..

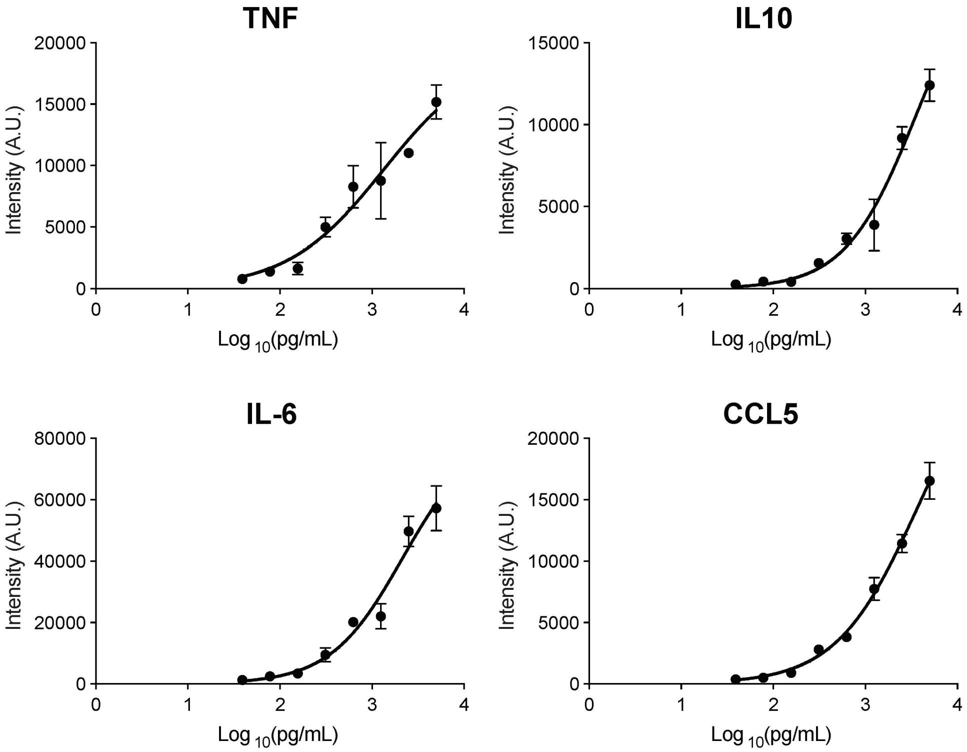


**Figure S4 (related to Fig. 4) Recombinant protein standard curves to convert fluorescent intensities measured from the microwell assay to protein concentrations.** Recombinant proteins were incubated with flow-patterned antibodies at a range of concentrations from (39-5000 pg/mL), and then were analyzed for intensity according to the same method as the microwell assay. The resulting standard curves were fit to a 4-parameter logistic curve. Recombinant protein data are mean ± SEM pooled from two biological replicates to create a consensus standard curve.

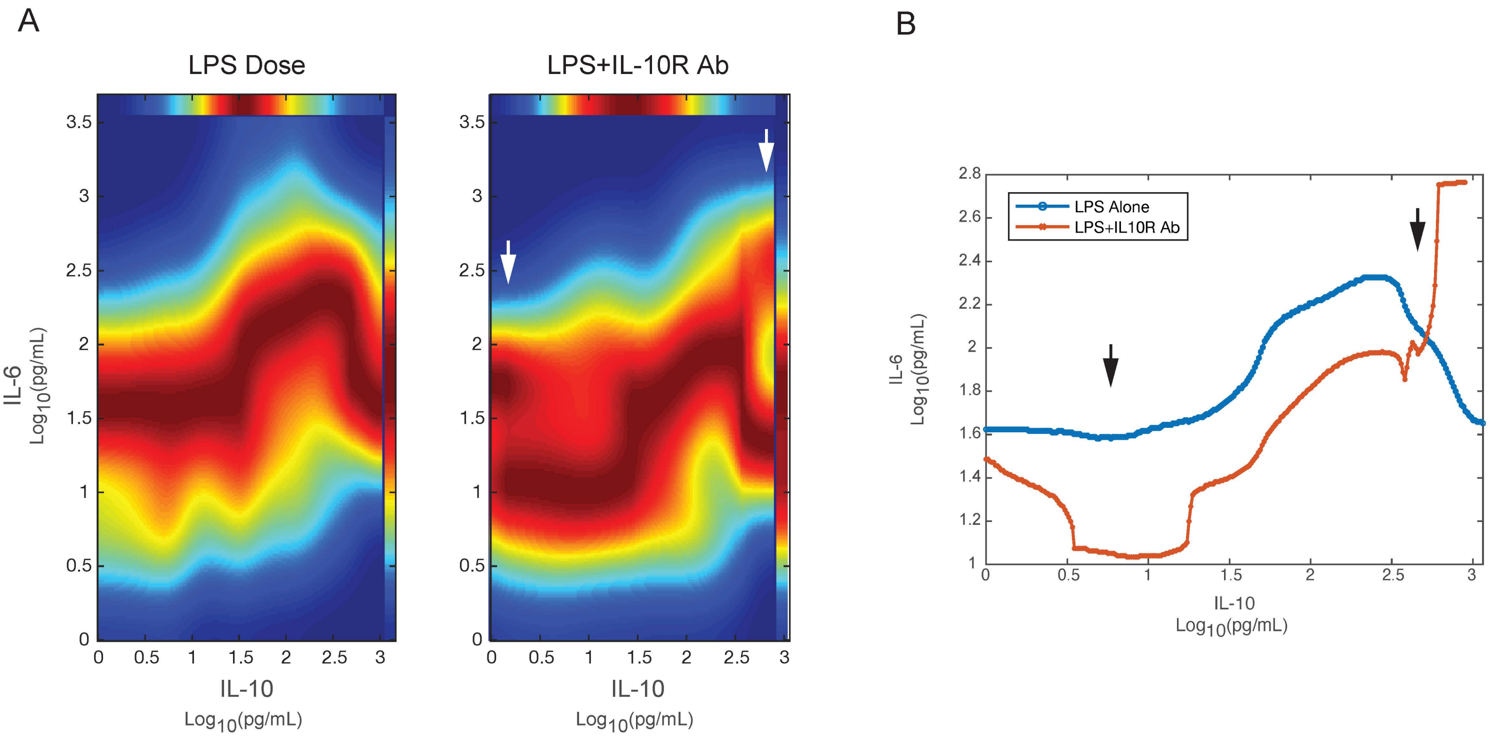


**Figure S5 (related to Fig. 4) Low levels of IL-10 prevent proinflammatory activation of low and non-responder cells. High levels of IL-10 diminish the proinflammatory response of high-responder cells.** (A) Conditional density re-scaled visualization (DREVI) plots showing the relationship between IL-10 and IL-6. Analysis was conducted on interpolated concentration data from fluorescent intensities in the microwell device. Data shown are pooled to include all stimulation doses of LPS (0, 10, 100, 1000 ng/mL) with or without IL-10R Ab at 30 µg/mL, where the 2 indicated cytokines were co-secreted during the 8-hour incubation (LPS alone: n=362, +IL-10R Ab: n=181). Arrows indicate areas of low and high IL-10 secretion most affected by IL-10 negative feedback. (B) Edge response functions were fit according to the conditional mean at the region of highest conditional density in the DREVI plots shown in A.

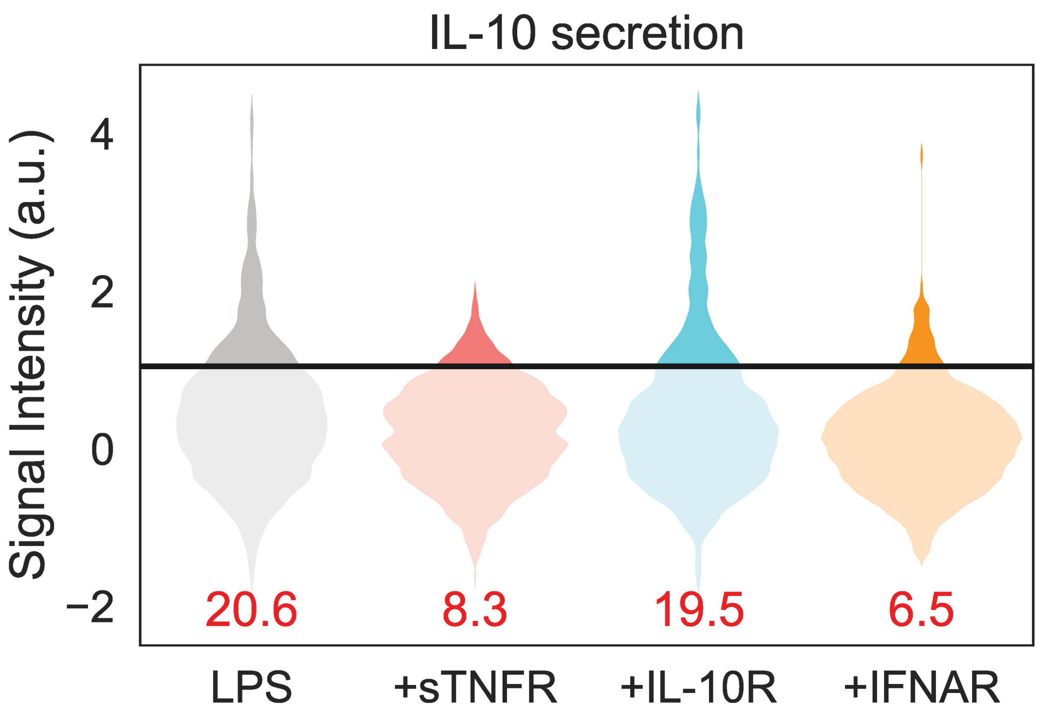


**Figure S6 (related to Fig. 5) Blocking TNF and IFN-β autocrine signaling suppresses IL-10 secretion in TLR4-stimulated BMDMs in the microwell device.** Violin plots of IL-10 secretion from individual BMDMs stimulated with 100 ng/mL LPS alone or co-stimulated with sTNFR, IL-10R, or IFNAR as indicated in Fig. 5 for 8 hours in the microwell assay. Data include only responder cell population (secreting at least one protein from Table 1). Black bar indicates fluorescent threshold of detection. Numbers in red indicate percentage of responder cells producing IL-10 above threshold (excludes non-responders).

**Supplemental Experimental Procedures**

**Flow Cytometry**

Plated BMDMs were stimulated for 15 minutes with LPS at varying doses (0, 0.1, 1, 10, 100, 1000 ng/mL). Afterwards, BMDMs were lifted in ice-cold PBS+EDTA with gentle scraping and immediately fixed with Phosflow Fix buffer (BD Biosciences) for 10 minutes at 37° C. Cells were then transferred to a 96 well u-bottom plate. After washing, cells were permeabilized with Phosflow Perm Buffer III (BD Biosciences). For intracellular phospho-protein staining, cells were blocked with Fc receptor antibody (eBioscience, CD16/32, 1:200 dilution) on ice for 15 mins in FACS buffer (PBS + 2% FBS). Cells were then stained in 50 µL using anti-Phospho-p38 MAPK (Thr180/Tyr182) (3D7) Rabbit mAb (Alexa Fluor 488 Conjugate) (Cell Signaling Technology #41768, 1:50 dilution). All samples were acquired on an Attune NxT Flow Cytometer, and analyzed with FlowJo (FlowJO, LLC).
